# Genetic polymorphisms in *COMT* and *BDNF* influence synchronization dynamics of human neuronal oscillations

**DOI:** 10.1101/2021.11.16.468778

**Authors:** Jaana Simola, Felix Siebenhühner, Vladislav Myrov, Katri Kantojärvi, Tiina Paunio, J. Matias Palva, Elvira Brattico, Satu Palva

## Abstract

Neuronal oscillations, their inter-areal synchronization, and scale-free dynamics constitute fundamental mechanisms for cognition by regulating communication in neuronal networks. These oscillatory dynamics have large inter-individual variability that is partly heritable. However, the genetic underpinnings of oscillatory dynamics have remained poorly understood. We recorded resting-state magnetoencephalography (MEG) from 82 healthy participants and investigated whether oscillation dynamics were influenced by genetic polymorphisms in Catechol-O-methyltransferase (*COMT*) Val^158^Met and brain-derived neurotrophic factor (*BDNF*) Val^66^Met. Both *COMT* and *BDNF* polymorphisms influenced local oscillation amplitudes and their long-range temporal correlations (LRTCs), while only *BDNF* polymorphism affected the strength of large-scale synchronization. Our findings demonstrate that *COMT* and *BDNF* genetic polymorphisms contribute to inter-individual variability in local and large-scale synchronization dynamics of neuronal oscillations. Comparison of these results to computational modelling of near-critical synchronization dynamics further suggested that *COMT* and *BDNF* polymorphisms influenced local oscillations by modulating the excitation-inhibition balance according to the brain criticality framework.

## Introduction

Neuronal oscillations and their inter-areal synchronization play fundamental mechanistic roles in cognitive functions, regulating neuronal processing and communication across distributed brain areas ^1–3^. In humans, dynamics of neuronal oscillations and their long-range phase synchronization, observed non-invasively in magneto- (MEG) and electroencephalography (EEG) recordings, serve fundamental roles in a variety of sensory, motor, and cognitive functions ^4–11^. During rest, brain activity involves frequency-specific inter-areal correlation structures throughout the 1–100 Hz range ^12–19^. The intrinsic spatial organization of spontaneous resting- state functional connectivity (FC) observed with MEG ^15, 17, 20^, intra-cranial EEG ^18^ and functional magnetic resonance imaging (fMRI) ^21, 22^ predicts individual variability in task performance. Resting-state FC networks reflect individual trait-like behavior ^23^ that could arise from the underlying individual brain structural organization ^24, 25^ and neuromodulation ^26–29^.

Amplitude envelopes of fast (> 1 Hz) oscillations are also characterized by slow (0.1–1 Hz) and infra-slow (0.01–0.1 Hz) fluctuations ^12, 30, 31^. These fluctuations are scale-free and exhibit power-law autocorrelations in their strength known as long-range-temporal correlations (LRTCs) ^30, 32–36^. In addition to instantaneous oscillations dynamics, inter-individual variability of LRTCs also predicts variability in task-related behavioral LRCTs ^36, 37^. Scale-free LRTCs are hallmarks of systems with critical or near-critical dynamics, suggesting that the brain operates near a critical phase transition between disorder and order, *i.e.*, between inadequate and excessive synchronization, respectively ^38^, which enables high flexibility and variability.

A large fraction of the variability in fast and slow oscillation dynamics could be heritable. MEG spectral power shows similarities between siblings ^39, 40^. Twin studies indicate that frontal midline theta activity is genetically related to symptoms and response time variability in Attention-Deficit/Hyperactivity Disorder (ADHD) ^41^ and that peak frequencies ^42^ as well as LRTCs ^43^ of local cortical oscillations have a genetic basis, so that up to 60% of the variance in neuronal LRTCs ^43^ and up to 85% of the variance in cognitive ability ^44^ can be attributed to genetic factors. However, what these genetic factors are and how they could contribute to the inter-individual variability of synchronization dynamics, including oscillation amplitudes, large- scale synchronization, and LRTCs, have remained central unresolved questions.

We hypothesized that polymorphism in genes regulating neuromodulation is one genetic factor underlying variability in oscillatory dynamics. Compelling evidence indicates that especially the neuromodulatory effects of the catecholamines dopamine (DA) and noradrenaline (NE) ^27, 28, 45, 46^ and the indolamine serotonin (5-hydroxytryptamine; 5-HT) ^47–49^ contribute to variability in neuronal dynamics ^26, 49–51^.

Brain DA and NE levels are regulated by their catabolism via COMT enzyme ^52, 53^ whose activity is influenced by a common single-nucleotide polymorphism (SNP) in the *COMT* gene substituting methionine (Met) for valine (Val) at codon 158 at sequence 4680 (Val^158^Met (rs4680)). *COMT* Val^158^ homozygotes have higher COMT activity and therefore lower brain DA and NE levels than Met^158^ homozygotes, with activity for heterozygotes falling in the middle ^54^. A considerable fraction of the variance in serotonin levels, on the other hand, is explained by Val^66^Met (rs6265) polymorphism of the gene coding for BDNF, which influences 5-HT receptor binding ^55^ and the activity-dependent secretion of BDNF ^56^. Convincing evidence emerging repeatedly from numerous fMRI studies indicates that *COMT* Val^158^Met (rs4680) polymorphism is associated with modulation of signal especially in the prefrontal cortex (PFC) and with cognitive functions dependent on PFC ^46, 57–60^ while *BDNF* Val^66^Met polymorphism is implicated in in many aspects of behavior and cognition ^61–63^. In addition to modulation 5-HT levels, activity- dependent release of BDNF also influences the maintenance and formation of neuronal networks ^64, 65^, thereby influencing cognition and behavior over lifetime.

Although the effects of *COMT* Val^158^Met and *BDNF* Val^66^Met polymorphisms on slow neuronal dynamics have been well-studied well with fMRI, their influence of fast neuronal oscillations has remained uncharted. We hypothesized that *COMT* Val^158^Met and *BDNF*Val^66^Met polymorphisms could influence inter-individual variability in oscillation amplitudes, phase synchronization, and their LRTCs. Importantly, critical dynamics in the brain is thought to be controlled primarily by the excitation-inhibition (E/I) ratio so that critical dynamics emerge at balanced E/I while excess of inhibition or excitation leads to sub- or super-critical dynamics, respectively ^34, 66, 67^. We thus further hypothesized that these polymorphisms would specifically exert their influence on critical brain dynamics via changes in the E/I.

Genome-wide association (GWAS) studies have demonstrated the clinical relevance of genes on functional and structural MRI data ^68^ and for local oscillatory activity using sensor-level EEG ^69^. Since our specific interest here was on the neuromodulation related genetic influences of polymorphisms in *COMT* Val^158^Met and *BDNF* Val^66^Met candidate genes on neuronal oscillations, we did not use a GWAS approach (which would require a huge sample size N > 1000) nor polygenic risk scores (which do not exist for oscillation dynamics).

To test our hypotheses, we measured resting-state brain dynamics with MEG and estimated local oscillation amplitudes, large-scale phase synchronization, and LRTCs from individually source-reconstructed MEG data. We then investigated their correlation with the *COMT* Val^158^Met and *BDNF* Val^66^Met polymorphisms estimated from SNP analysis (Figure 1) and compared the results with predictions from computational modeling. In line with the predictions of the brain criticality framework and computational modeling, we found that the *COMT* Val^158^Met and *BDNF* Val^66^Met polymorphisms influenced local oscillation amplitudes and LRTCs, while *BDNF* Val^66^Met polymorphism also influenced the strength of large-scale synchronization.

**Figure 1.**
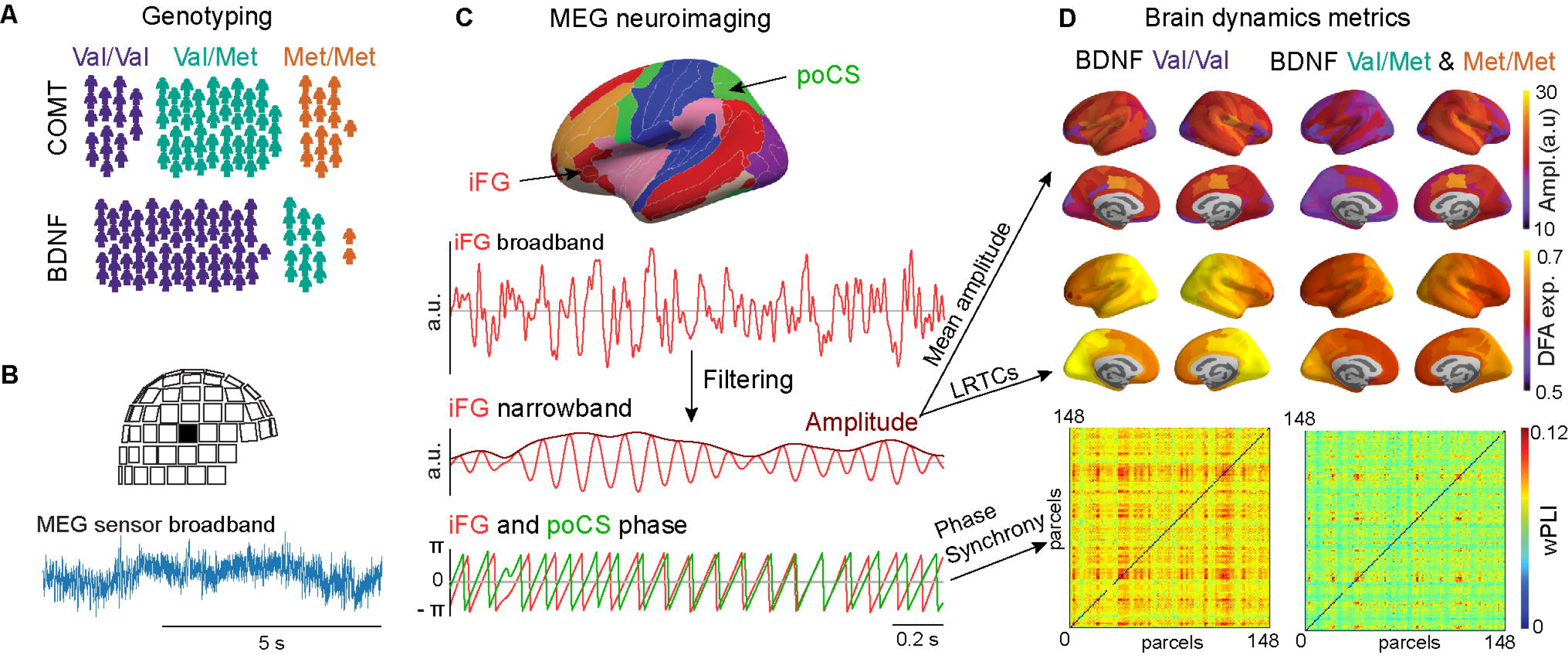
A schematic of study outline and rationale. (A) Genotyping of the participants (*N* = 82) was used to identify the *COMT* Val^158^Met (rs4680) and *BNDF* Val^66^Met (rs6265) genetic polymorphisms. (B) Broadband MEG data were recorded from each participant with 306-channel MEG during ∼8-minute resting state. (C) MEG data were source-reconstructed using minimum-norm estimation with individual MRI-based head and cortical surface anatomy and collapsed to cortical parcels (upper panel). Parcel-wise broadband signals (shown for inferior frontal gyrus, iFG, in 2^nd^ panel) were filtered to narrowband signals (signal and amplitude envelope (shown here for the iFG in 3^rd^ panel). From the narrowband signal, instantaneous phase was extracted (shown for iFG and posterior central sulcus, poSC, in lower panel). Mean amplitudes and LRTCs were estimated for each parcel from amplitude envelope time series, and inter- parcel phase synchronization for all parcel pairs from phase time series. (D) In group analyses, we assessed the anatomical distribution of mean amplitudes, LRTCs, and synchronization within the polymorphism groups and identified statistically significant differences between these groups (Figures 2-4).

## Results

### *COMT* Val/Met and *BDNF* Val polymorphisms are associated with the strongest oscillation amplitudes

We first evaluated mean oscillation amplitudes between 3–60 Hz averaged across cortical parcels separately for participants grouped according to the *COMT* Val^158^Met (Val/Val, Val/Met, Met/Met) and the *BDNF* Val^66^Met (Val/Val and the combined Val/Met and Met/Met) polymorphisms. As observed commonly in resting state, oscillation amplitudes peaked in the alpha band (α, 8–14 Hz) (Figure 2A). We then clustered oscillations to frequency bands based on the spatial similarity using Louvain clustering (Figure S1) that revealed similarity in theta (θ, 3–7 Hz), alpha, beta (β, 14–30 Hz) and gamma (γ, 30–60 Hz) frequency bands. Repeated measures ANOVA revealed differences in the oscillation strength between these canonical bands (Figure S2). Using univariate ANOVAs that controlled for age effects, we further showed both in the wavelet frequencies (Figure 2A) and in the canonical frequency bands (Figure S2) that *COMT* Val^158^Met and *BDNF* Val^66^Met polymorphisms had no impact on oscillation amplitudes at the whole-brain level. We further confirmed that age or the gender ratios did not differ between the polymorphism groups (Tables 1 and 2).

**Figure 2.**
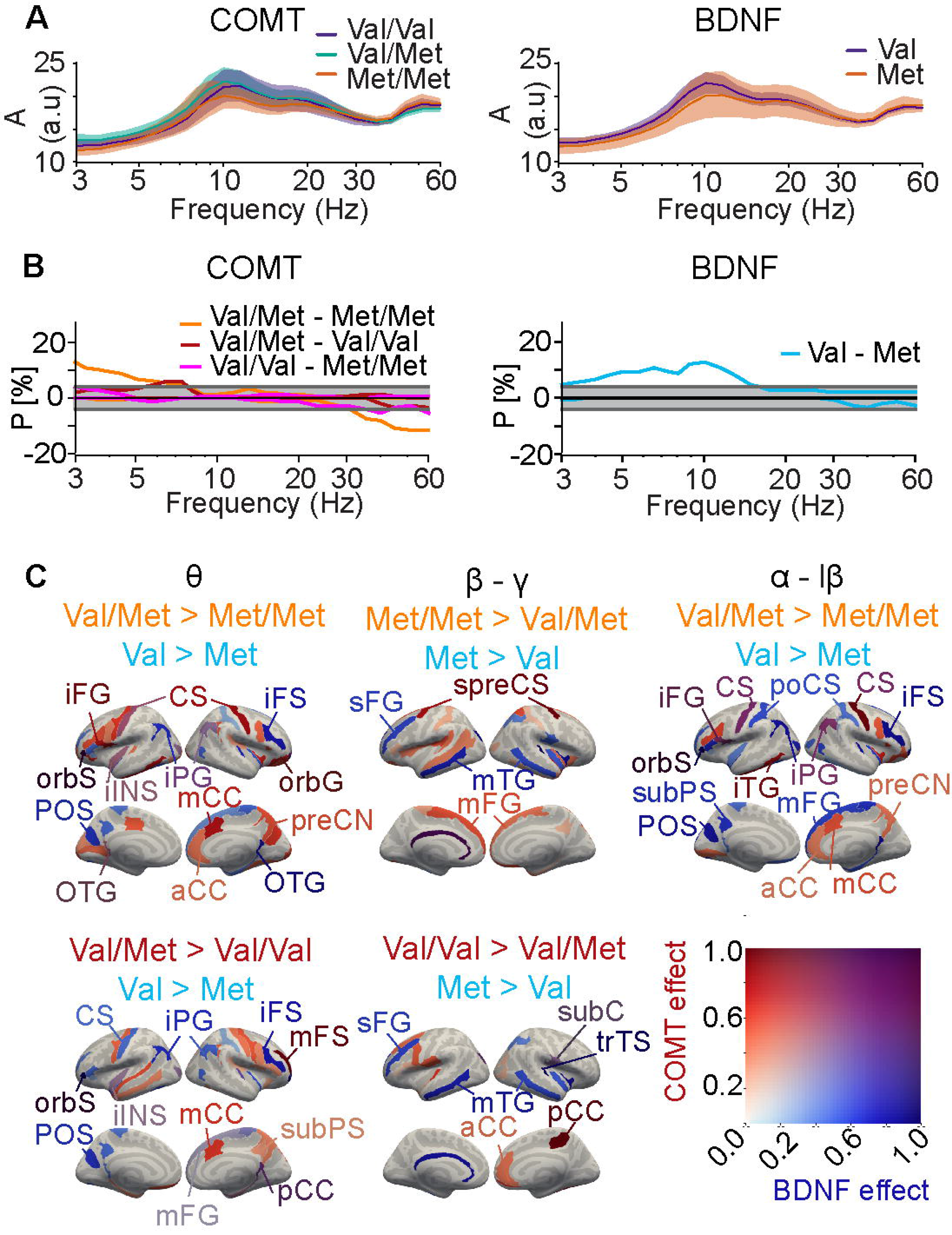
*COMT* and *BDNF* polymorphisms influence oscillation amplitudes. (A) Oscillation amplitudes averaged across parcels for the *COMT* (Val/Val, purple, *n* = 18; Val/Met, turquoise; *n* = 48; and Met^/^Met, orange, *n* = 16) and *BDNF* polymorphism groups (Val/Val, purple, *n* = 66; Met carriers, orange, *n* = 16). Shaded areas represent 95% bootstrapped confidence intervals of the group means. (B) The fraction *P* of parcels with a significant positive and negative pairwise difference (shown separately) between the *COMT* and *BDNF* polymorphisms groups in oscillation amplitudes (Mann–Whitney-U test, *p* < 0.05). The light grey area indicates the 5% alpha level for findings expected by chance under the null hypothesis and the dark grey area indicates the range (3% total) in fractions of significant parcels observable by chance at *p_q_* < 0.05 in any single frequency band across the spectrum (26 frequencies). (C) Cortical parcels where the amplitudes differed significantly between polymorphism groups (indicated above the brain surfaces) in the θ (3–7 Hz), β–γ (20–60 Hz), and α to low-β (7–18 Hz) frequency ranges in at least one of the individual frequencies. The color scale indicates the max-normalized amplitude differences for *COMT* (red) and *BDNF* (blue) polymorphisms independently, so that parcels where the effects are co-localized have purple hues. Abbreviations are formed from a combination of i) Areas: a = anterior, ca= calcarine, i = inferior, m = middle, p = posterior, s = superior, tr = triangular and ii) Lobes: C= central / cingulate, F = frontal, P = parietal, O = occipital, T = temporal, and iii) G = Gyrus, S = Sulcus. Additional abbreviations: subC = subcentral gyrus/sulcus, preCN = precuneus.

**Table 1.** Mean sample characteristics (SD in parentheses) by the COMT polymorphisms.

|  | Val/Val | Val/Met | Met/Met |
| --- | --- | --- | --- |
| N | 18 | 48 | 16 |
| Age mean, years <sup>1</sup> | 30.65 (9.90) | 28.53 (8.91) | 29.87 (10.69) |
| Age range | 22–55 | 19–50 | 18–53 |
| Handedness (right/left) <sup>2</sup> | 18/0 | 41/5 | 15/1 |
| Gender (Female/Male) <sup>3</sup> | 11/7 | 27/21 | 6/10 |
| Rs mean, minutes <sup>4</sup> | 7.86 (2.20) | 7.97 (3.28) | 7.49 (2.69) |
| Rs range | 5–10 | 5–24 | 5–12 |

<sup>1</sup> Age information missing from three participants. The polymorphism groups did not differ in terms of age as confirmed by a one-way ANOVA.
<sup>2</sup> Handedness information missing from two participants.
<sup>3</sup> The percentages of female and male participants did not differ within the *COMT* polymorphism groups as indicated by the Chi-squared test.
<sup>4</sup> Rs = resting state. The duration of the resting state recording did not differ between polymorphism groups as confirmed by a one-way ANOVA.

**Table 2.**
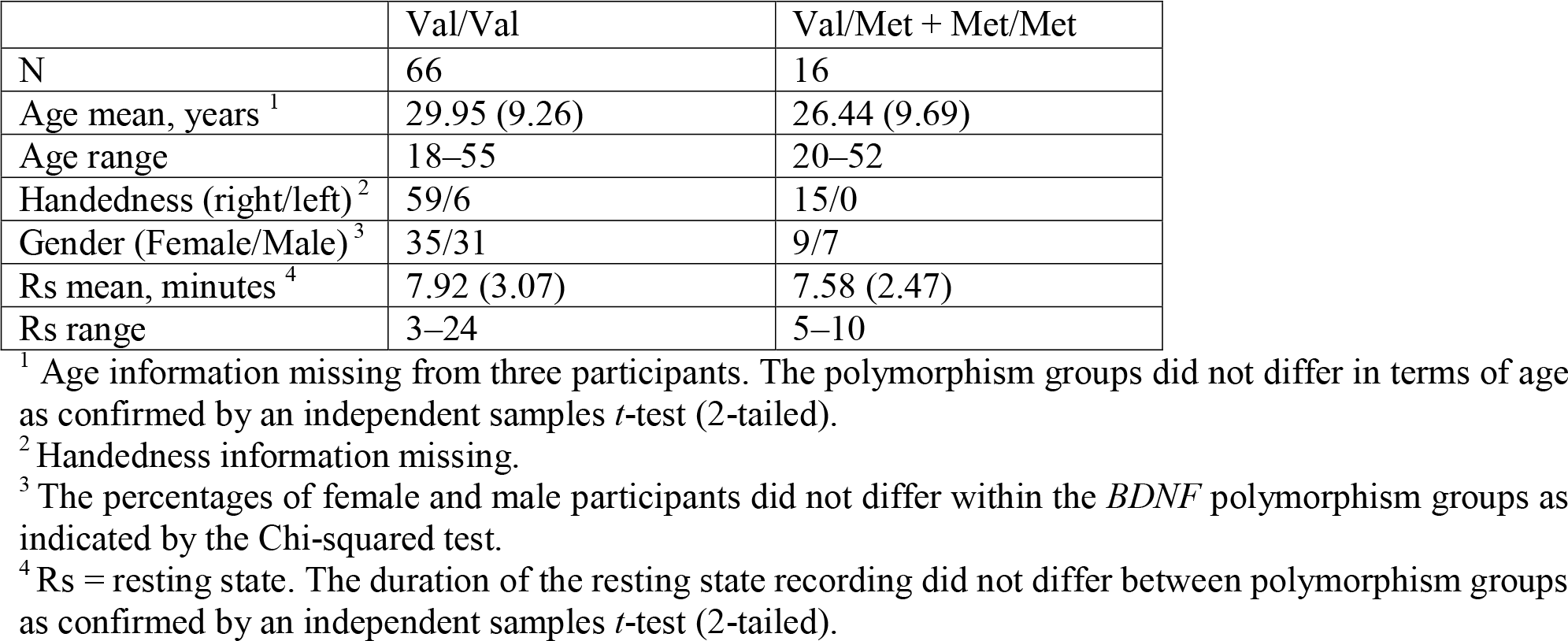
Mean sample characteristics (SD in parenthesis) by the *BDNF* polymorphisms.

|  | Val/Val | Val/Met + Met/Met |
| --- | --- | --- |
| N | 66 | 16 |
| Age mean, years <sup>1</sup> | 29.95 (9.26) | 26.44 (9.69) |
| Age range | 18–55 | 20–52 |
| Handedness (right/left) <sup>2</sup> | 59/6 | 15/0 |
| Gender (Female/Male) <sup>3</sup> | 35/31 | 9/7 |
| Rs mean, minutes <sup>4</sup> | 7.92 (3.07) | 7.58 (2.47) |
| Rs range | 3–24 | 5–10 |

<sup>1</sup> Age information missing from three participants. The polymorphism groups did not differ in terms of age as confirmed by an independent samples *t*-test (2-tailed).
<sup>2</sup> Handedness information missing.
<sup>3</sup> The percentages of female and male participants did not differ within the *BDNF* polymorphism groups as indicated by the Chi-squared test.
<sup>4</sup> Rs = resting state. The duration of the resting state recording did not differ between polymorphism groups as confirmed by an independent samples *t*-test (2-tailed).

To identify possible anatomically limited polymorphism effects, we tested significant pairwise differences between polymorphism groups for individual parcels and for each frequency. For *COMT*, the amplitudes were stronger (Mann-Whitney-U test, *p* < 0.05) for Val/Met heterozygotes than for either Met or Val homozygotes in the θ band (3–7 Hz) (in up to 13% and 6% of cortical parcels, respectively) (Figure 2B), whereas in the high-β to γ (20–60 Hz) bands, amplitudes were larger for Met homozygotes (12% of parcels compared to Val/Met, 6% compared to Val/Val). For *BDNF*, oscillation amplitudes were stronger for the Val homozygotes than for Met carriers in θ and in the α to low-β bands (up to 13% of parcels). We next visualized significant differences in oscillation amplitudes between the polymorphism groups in the cortical anatomy (Figure 2C). Stronger θ and α−low-β band oscillation amplitudes for the *COMT* Val/Met heterozygotes were localized to the central sulcus (CS), anterior, mid- and posterior cingulate cortex (aCC, mCC, pCC), inferior frontal gyrus (iFG), right-hemispheric middle frontal gyrus (mFG) and inferior parietal gyrus (iPG), inferior part of the insula (iINS) and precuneus (preCN) (Figure 2C) these parcels mostly belonging to the frontoparietal network (FPN) and default mode networks (DMN) (Figure S3). Of these, iPG and left iINS and left CS showed also larger amplitudes for *BDNF* Val homozygotes than for Met carriers. In the β-γ bands, larger amplitudes for *COMT* Met homozygotes compared to Val/Met heterozygotes were observed in superior precentral sulcus (spreCS), corresponding to frontal eye fields (FEF) and in temporal cortices while stronger amplitudes for the Val homozygotes were found in aCC and pCC. For *BDNF* Val homozygotes, oscillation amplitudes were larger than in Met carriers in iFS, parieto-occipital sulcus (POS) and postcentral sulcus (poCS) in θ to low-β bands. *COMT* and *BDNF* jointly affected θ and α to low-β band oscillation amplitudes in the CS and iPG.

### *COMT* Val^158^Met and *BDNF* Val^66^Met polymorphisms influence DFA scaling exponents

We quantified long-range temporal correlations (LRTCs) in oscillation amplitude fluctuations with detrended fluctuation analysis (DFA) ^70^ and estimated the mean DFA scaling exponents in all wavelet frequencies (3–60 Hz). DFA exponents were first averaged across parcels separately for each *COMT* Val^158^Met and *BDNF* Val^66^Met polymorphism group for each frequency (Figure 3A). Repeated measures ANOVA revealed differences in the DFA exponents between canonical frequency bands (Figure S2). Univariate ANOVAs that controlled for age showed that *COMT* polymorphism had an effect on whole-brain DFA exponents in the γ frequency band averaged across the brain [*F*(2, 75) = 3.80, *p* = .027, η ^2^ = 0.092] (*n* = 79) (Figure 3A and S2). Group mean exponents were higher at the γ band for *COMT* Val/Met individuals than for Val/Val (*t*(64) = 2.08, *p* = 0.043, uncorrected) and Met/Met (*t*(62) = 3.50, *p* = 0.001) individuals when equal variances were not assumed. The *BDNF* polymorphism affected whole-brain DFA exponents in the canonical α band [*F*(1, 76) = 4.19, *p* = .044, η ^2^ = .052], when age was controlled for (*n* = 79), with Val homozygotes exhibiting larger LRTC exponents than Met carriers (Figure 3A and S2).

**Figure 3.**
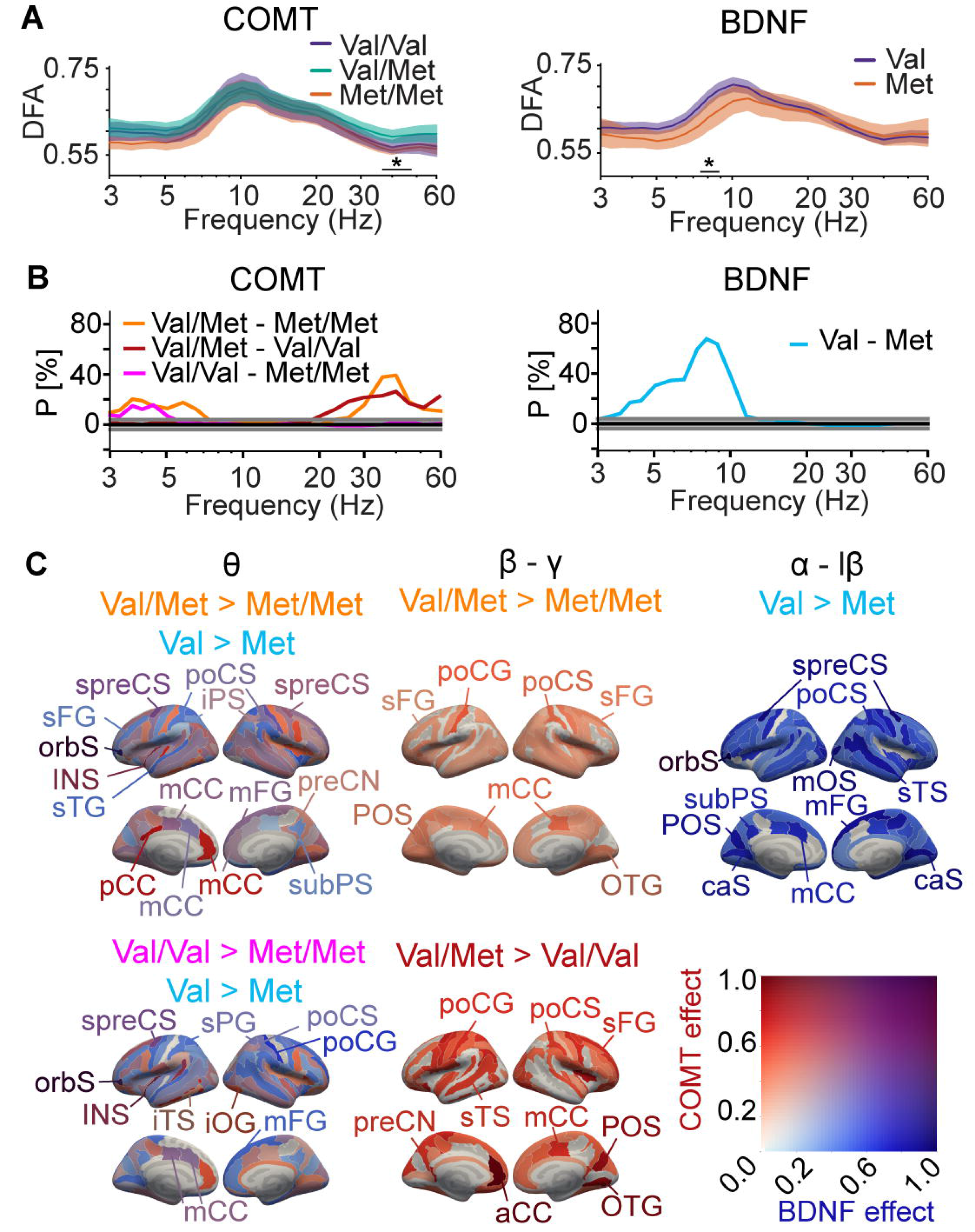
Influence of *COMT* and *BDNF* polymorphisms on LRTCs. (A) Detrended fluctuation analysis (DFA) exponents averaged across parcels for the *COMT* and *BDNF* polymorphism groups. Shaded areas represent 95% bootstrapped confidence intervals of group means. Black bar with asterisk denotes the range of frequencies with significant differences between the *COMT* and *BDNF* polymorphism groups (univariate ANOVA, with age as covariate, *: *p* < 0.05, uncorrected). (B) Fractions (*P*) of parcels with a significant positive and negative difference between C*OMT* and *BDNF* polymorphism groups in the DFA exponents (Mann–Whitney-U test, *p* < 0.05), as in Figure 2B. (C) Cortical areas where significant differences between polymorphism groups in the DFA exponents were found in the θ, β–γ and α– low-β bands (Mann–Whitney U test, *p* < 0.05, corrected). Colors and abbreviations as in Figure 2C.

Similar to oscillation amplitudes, *COMT* Val^158^Met and *BDNF* Val^66^Met polymorphisms had a more robust effect on DFA exponents at the resolution of cortical parcels indicating large anatomical heterogeneity. DFA exponents were greater for *COMT* Val/Met heterozygotes than for Met homozygotes (Mann-Whitney-U test, *p* < 0.05) in the θ band (up to 20% of parcels) and in the β–γ band (up to 40%) (Figure 3B). Further, θ band DFA exponents were greater for Val/Val than for Met/Met (16%) and β–γ DFA exponents were greater in Val/Met than for Val/Val (26%) (Figure 3B). For *BDNF,* a very robust effect of polymorphism was found for DFA exponents.

These were greater for Val homozygotes than Met carriers in the θ band (35%) and in α band (68%) (Figure 3B). The localization of significant differences in DFA exponents between polymorphism groups (Mann-Whitney-U test, *p* < 0.05, Figure 3C) revealed wide-spread group differences. In the θ band, large differences between *COMT* Val/Met heterozygotes and Met homozygotes and between *BDNF* Val homozygotes and Met carriers were co-localized to preCS, mCC, and mFG (Figure 3C). DFA exponents in θ band were also greater for *COMT* Val/Met group in INS, pCC, and in β–γ band particularly in parcels belonging to somatomotor (SM) network and DMN. *BDNF* Val homozygotes had greater LRTC exponents in α and low-β bands than Met carriers in nearly all parcels (Figure 3C). Generally, the effects of *COMT* and *BDNF* polymorphisms on LRCTs were widespread, in line with prior work and with the idea that LRTCs reflect global fluctuations in neuronal activity across the brain’s modular structure ^16, 30, 36^.

Finally, to gain insight for the common effects of *COMT* Val^158^Met and *BDNF* Val^66^Met polymorphisms on oscillation amplitudes and DFA exponents, we co-localized their effects. We found that joint effects on amplitudes and DFA exponents were strongest in the parcels of PFC and PPC, CS and anterior cingulate structures (Figure S4).

### *BDNF* polymorphism correlates with the strength of large-scale phase-synchronization

We next set out to assess the influence of *COMT* Val^158^Met and *BDNF* Val^66^Met polymorphisms on inter-areal phase synchronization of neuronal oscillations. Phase synchrony is a key characteristic of individual cortical dynamics and has fundamental influence on behavior ^1, 3, 4, 7, 14, 22^. We estimated phase synchrony among all cortical parcels with the weighted phase-lag index (wPLI) that is insensitive to linear source mixing ^71^ and computed the mean synchronization for each of the *COMT* Val^158^Met and *BDNF* Val^66^Met polymorphism groups (Figure 4A). We did not find significant differences (Kruskal-Wallis test, *p* < 0.05) among the *COMT* Val^158^Met polymorphism groups. Mean synchronization for *BDNF* Val homozygotes was stronger than for Met carriers in the θ and α frequencies (Figure 4A).

**Figure 4.**
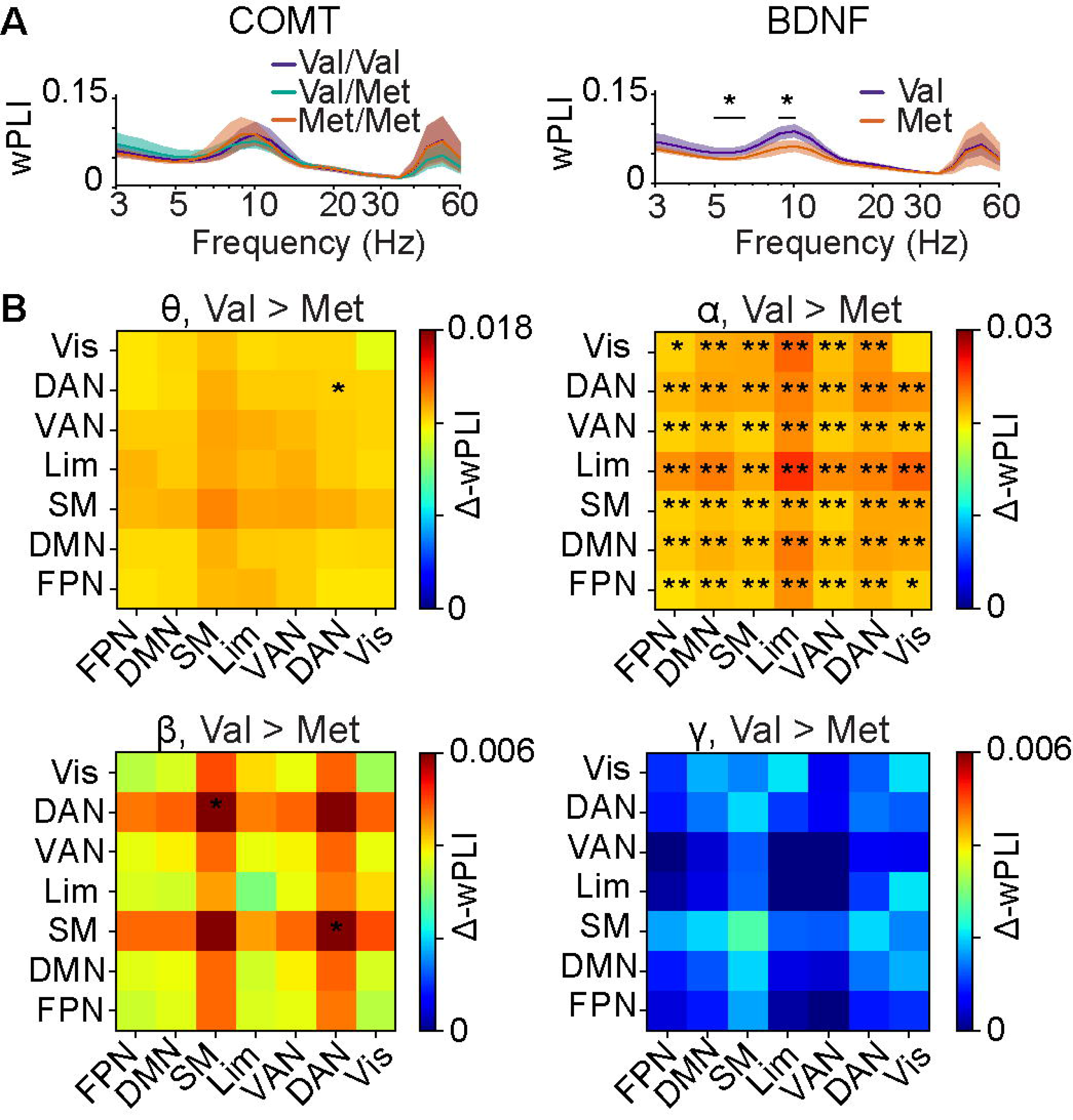
*BDNF* polymorphisms influence large-scale phase-synchronization. (A) The mean strength of phase-synchronization averaged across the whole brain for the *COMT* Val^158^Met and *BDNF* Val^66^Met polymorphism groups with 95% bootstrapped confidence intervals. Black bars with asterisks denote the frequencies where a significant group difference was found (Kruskal-Wallis, *: *p* < 0.05, uncorrected). (B) The differences in the mean phase synchrony (⊗-wPLI) between *BDNF* Val homozygotes and Met carriers within and between functional systems, averaged across hemispheres and wavelet frequencies in canonical frequency bands. Red color indicates stronger synchronization for *BDNF* Val^66^ homozygotes and blue for Met carriers. Stars denote the brain system pairs where the group difference was significant (Kruskal-Wallis, *: *p* < 0.05, uncorrected, **: *p* < 0.01, Benjamini-Hochberg corrected). Abbreviations: DAN = Dorsal attention network, DMN = Default Mode network, FPN = Frontoparietal network, Lim = Limbic system, SM= Somatomotor network, VAN = Ventral attention network, Vis = Visual network.

To acquire neuroanatomical insight into the patterns of inter-areal synchronization in functional brain systems, we collapsed the parcel-wise synchronization estimates within functional systems ^72^ (Figure S3) and canonical frequency bands (Figure S1). Synchronization between or within the subsystems did not differ significantly among the *COMT* Val^158^Met polymorphisms (Figure S5). However, *BDNF* Val homozygotes exhibited stronger synchronization than Met carriers in the α band between and within nearly all subsystems (Figure 4B and Figure S5, Kruskal-Wallis test, corrected with Benjamini-Hochberg). We also computed synchronization separately for each hemisphere and between hemispheres. Interestingly, group differences in α band synchronization were widespread and more robust for left- and inter- hemispheric connections than for the right-hemispheric connections (Figure S6). Additionally, β- band synchronization was stronger for *BDNF* Val homozygotes than Met carriers within and between several left-hemispheric networks, including FPN, dorsal attention network (DAN), ventral attention (VAN) and Limbic networks (Lim) and DMN (Figure S6).

### Computational model-based approach to neuromodulation effects on critical synchronization dynamics

We next set out to address the effects of *COMT* Val^158^Met and *BDNF*Val^66^Met polymorphisms in the context of brain criticality. Brain criticality is assumed to be regulated by a control parameter (K) that reflects the gross excitation / inhibition (E/I) ratio ^66, 73, 74^. The human brain is thought to normally operate in a slightly subcritical regime ^75^, where LRTCs exhibit large inter-individual variability ^32, 33, 36, 37^. We hypothesized that *COMT* Val^158^Met and *BDNF* Val^66^Met polymorphisms would bias the E/I balance and hence the “control parameter” that tunes the distance to criticality.

### The carriers of different variants should therefore be at different positions away from the critical point

To assess the relationships of the observable brain dynamics measures with the control parameter, we implemented a nested Kuramoto model with separable local network (cortical parcel) and large-scale (whole-brain network) levels ^15^ of synchronization dynamics exhibiting a transition from sub- to super-critical dynamics (Figure 5A). We used the model to predict synchronization dynamics of neuronal oscillations for two hypothetical cohorts (A and B) that differed in values of the control parameter (K) regulating their position in the critical regime. In the model, increasing K led to parallel strengthening of local amplitudes, large-scale synchronization, and LRTCs throughout the subcritical regime up to the critical point (Figure 5A) so that the cohort B with larger K exhibited larger oscillation amplitudes, LRTCs, and large-scale synchronization (Figure 5A-B), and as a novel model-predicted measure, larger mutual correlations (Figure 5C) between these three values.

**Figure 5.**
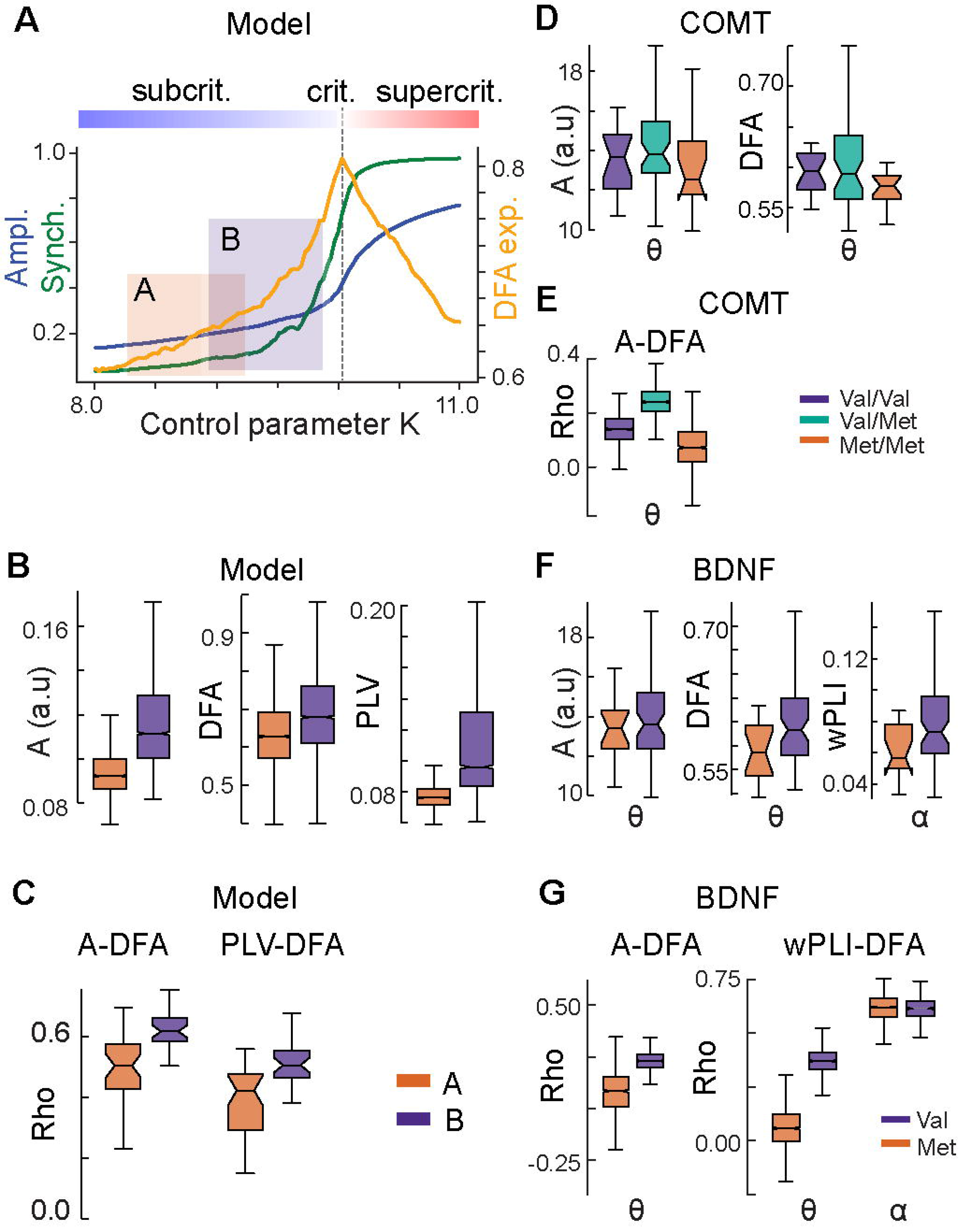
Model and experimental evidence for modulation of brain criticality. (A) Computational model of local and inter-areal synchronization dynamics in the realistic human connectome shows that both local amplitude (blue) and inter-areal synchronization (green) increase monotonically with increasing control parameter K and exhibit a phase-transition between low- (subcritical) and high-synchronization (supercritical) regimes. Conversely, the DFA scaling exponent (yellow), a measure of long-range temporal correlations (LRTCs), peaks at the critical point. The model predicts that two cohorts (A, B) with subjects operating in different parts of the subcritical regime should exhibit group differences in amplitude, LRTCs, and inter-areal synchronization. (B) Local amplitude A, DFA exponents, and inter-areal phase synchronization (PLV) in modeled data for two cohorts A (orange) and B (purple). All metrics are greater for the cohort B that is designed to operate closer to the critical point (see panel A) than cohort A. The box ends indicate the lower quartile (Q1, 2^5t^h percentile) and upper quartile (Q3, 7^5t^h percentile) across cortical regions/parcels, notches indicate the median, and whiskers indicate the range of Q1 – 1.5 * IQR and Q3 + 1.5 * IQR, where IQR is the inter-quartile range (Q3-Q1). (C) Correlations between amplitude and DFA exponents, and between PLV and DFA, in modeled data are also larger for cohort B than the more subcritical cohort A. (D) θ band A and DFA exponents for *COMT* Val^158^Met polymorphism groups in MEG data and (E) their correlation. (F) θ band A and DFA exponents, and α band global phase synchronization for the *BDNF* polymorphism groups in MEG data. (G) Correlation between A and DFA exponents and between wPLI and DFA exponents in θ- band differs between *BDNF* Val^66^Met polymorphism groups, but correlation between wPLI and DFA in α band does not.

To investigate whether *COMT* and *BDNF* polymorphisms could influence synchronization dynamics via modulation of K, in line with the brain criticality framework, we computed these dynamics separately for each polymorphism group. In line with model predictions, the elevated θ- band amplitudes and LRTCs in *COMT* Val/Met group (Figure 5D) were paralleled with stronger correlation of local oscillation amplitudes and LRTCs (Figure 5E, Val/Met > Met/Met in 95% and Val/Met > Val/Val in 84% of permutations). For *BDNF,* elevated θ-band amplitudes and LRTCs in Val homozygotes (Figure 5F) were associated with stronger correlation of local oscillation amplitudes and LRTCs (Figure 5G, Val > Met in 92% of permutations, for γ bands see Figure S7). In contrast, elevated α-band global synchronization in *BDNF* Val/Val group (Figure 5F) was not associated with elevated correlation with LRTCs (Figure 5G). These findings thus provide converging evidence that *COMT* Val^158^Met and *BDNF* Val^66^Met polymorphisms impact local oscillation dynamics as predicted by the framework of brain criticality under the hypothesis that these polymorphisms influence the control parameter of cortical critical dynamics. In contrast, the most robust effects of *BDNF* on large-scale synchronization were not in line of the model predictions, suggesting that these were mediated mainly by other factors than the control parameter of brain criticality.

## Discussion

Inter-individual variability in spontaneous oscillatory dynamics influences individual behavioral performance and cognitive ability ^4, 15, 22, 76, 77^. While the causes of this variability are diverse, part of it is attributable to genetics, which has been shown for human MEG and EEG data ^39, 42, 43^. While it has been shown that macroscopic oscillations can be influenced through regulation of dynamic circuit motifs on a cellular level ^2, 39, 42^, there has been little research on how the genetic biases of cellular mechanisms translate into alterations of macroscopic brain dynamics observable in electrophysiological neuroimaging. We tested here a hypothesis that the *COMT* Val^158^Met and *BDNF* Val^66^Met genetic polymorphisms contribute to the variability in emergent local (oscillation amplitudes and LRTCs) and global (inter-areal phase synchronization) dynamics of neuronal oscillations. We showed that both *COMT* and *BDNF* polymorphisms influenced local oscillation amplitudes and LRTCs. However, only *BDNF* polymorphism also impacted the strength of global synchronization. Using computational modeling, we furthermore showed that the influence of *COMT* and *BDNF* on local oscillation dynamics is achieved in line with brain criticality framework. This suggests that *COMT* and *BNDF* polymorphisms influence synchronization dynamics via modulations of the brain E/I balance in line with the established role of *COMT* and *BDNF* in the regulation of neuronal activity via DA, NE, and 5-HT.

### Local oscillation dynamics are modulated by *COMT* and *BDNF* **polymorphisms**

Prior observations about *COMT* and *BDNF* polymorphism effects on electrophysiological activity are scarce and restricted to EEG and MEG sensor level ^78–80^. In these studies, *COMT* Val homozygotes were found to have lower α power than Met homozygotes ^79^, and *COMT* polymorphism was found to influence phase-resetting of γ oscillations in attention ^80^. We used here a data-driven approach for mapping the effects of *COMT* and *BDNF* polymorphism on both instantaneous oscillation amplitudes and their LRTCs from source-reconstructed MEG data. We found θ−α band oscillations amplitudes to be greater for *COMT* Val/Met heterozygotes than for *COMT* Val or Met homozygotes, and θ and α oscillation amplitudes to be greater for *BDNF* Val homozygotes than for Met carriers. In addition to local oscillation amplitudes, we also found that *COMT* and *BDNF* polymorphisms modulated LRTCs in oscillation amplitude dynamics and hence the variability of the slow fluctuations in oscillation strengths. To our knowledge, this study is the first to report the influence of genetic polymorphisms on LRTCs in oscillation amplitude fluctuations. LRTCs were stronger for *COMT* Val/Met heterozygotes than for Val or Met homozygotes in θ and γ bands and stronger for *BDNF* Val homozygotes than for Met carriers in α- low-β bands, demonstrating that near-critical dynamics of brain oscillations is influenced by *COMT* and *BDNF* polymorphisms.

Our results provide robust evidence for *COMT* polymorphism influencing local amplitude dynamics. As *COMT* Val/Met heterozygotes have intermediate COMT enzymatic activity, and therefore intermediate brain DA and NE levels compared to Val and Met homozygotes ^54^, and as operating near criticality has been proposed to be beneficial for individuals (see below), our results are consistent with previous findings showing that the impact of neuromodulation on cognition follows an “inverted-U-shape” function where both excessively low and excessively high neuromodulation impair performance ^49, 57, 59, 81–83^.

The effects of *COMT* Val^158^Met polymorphism were localized particularly to FPN and DMN, in agreement with the role of DA in cognitive control ^81, 84^, as well as with previous fMRI studies in which the association between the polymorphism and neuronal activity was predominantly observed in PFC ^46, 57–59^ which has low DA reuptake and higher COMT levels.

*BDNF* Val homozygotes exhibited stronger α band amplitudes bilaterally in frontoparietal regions and parcels of the visual system. In line with previous observations ^36^, differences in LRTC exponents were widespread, suggesting that they reflect state-dependent changes affecting most of the brain. Overall, these results support the idea that *COMT* and *BDNF* influence local oscillation dynamics in a variety of cortical regions important for cognition via DA, NE, and 5- HT.

### *BDNF* polymorphism influences whole-brain connectivity

We found that *BDNF* Val^66^Met but not *COMT* Val^158^Met polymorphism influenced large-scale global inter-areal phase synchronization. Global α band synchronization was stronger for *BDNF* Val homozygotes than Met carriers in all functional subsystems except within the visual system. *BDNF* Val homozygotes also had stronger θ synchronization within DAN and stronger left- hemispheric β synchronization between various subsystems, particularly involving DAN and DMN. This robust modulation of synchronization dynamics by *BDNF* polymorphism could contribute to significant differences across individuals in behavior and cognition, which have been shown to be strongly influenced by spatial and spectral characteristics of oscillatory synchronization ^15, 17, 24^. However, large-scale synchronization was not influenced by *COMT* Val^158^Met polymorphism, although it has been shown that neuropharmacologically elevated levels of catecholamines are related to fMRI-based changes in functional connectivity ^27^. These differences might be explained by different effects of catecholaminergic levels on BOLD signal and electrophysiological connectivity, or by *COMT* polymorphism having more subtle effects on neuronal dynamics compared to neuropharmacological interventions.

### *COMT* and *BDNF* influences oscillation via impact on brain critical dynamics

Near-critical synchronization dynamics has been proposed to enable transient and flexible computations that are beneficial for the individuals by maximizing information storage and transmission ^38, 75, 76, 85–88^. This notion has been supported experimentally by behavioral data ^89^ and *in vitro* studies ^66^. Here we show that genetic polymorphisms in *COMT* and *BDNF* influenced individual near-critical synchronization dynamics and proximity to the critical point. The systems-level E/I ratio (or gain) is thought to be the primary control parameter for the regulation of critical brain dynamics ^66, 74^. We postulated that the observed differences in neuronal synchronization dynamics are due to neuromodulation-related shifts in the system’s net E/I ratio. Comparison of experimental and computational-model data showed that *COMT* Val^158^Met and *BDNF* Val^66^Met polymorphisms influenced local oscillatory dynamics and their correlations in the θ band in line with the brain criticality framework. More specifically, *COMT* Val^158^Met heterozygotes and *BDNF* Val^66^ homozygotes had greater amplitudes and LRTCs and stronger correlations between LRTCs and oscillation amplitudes than the other polymorphism groups in θ band, indicating that they are associated with a higher value of the underlying control parameter, ., with greater net excitatory drive or neuronal gain ^51^. This is in accordance with the framework in which neuromodulatory drive influences whole-brain neural states ^90, 91^.

In contrast, we found no evidence that the influence of *BDNF* Val^66^Met polymorphism on global α-band synchronization could be explained within the brain criticality framework. This finding implies that the strongest effects of *BDNF* polymorphisms on global synchronization might not be due to a shifts in global excitation, but to other mechanisms such as the influence of BDNF on maintenance, maturation, and formation of neuronal networks ^64, 65^ putatively via receptor TrkB (neurotrophic receptor tyrosine kinase 2) signaling ^92, 93^. These could impact large- scale network synchronization via changes in structural connectivity ^24^ or via changes in brain gray matter whose density has been shown to be associated with neuroreceptor and neurotransporter availabilities ^94^ and whose thickness is associated with the structure of oscillatory networks ^95^.

## Conclusions

Our data showed that *COMT* Val^158^Met and *BDNF* Val^66^Met polymorphisms contribute to the variability of oscillation dynamics, their LRTCs, and synchronization dynamics, and adds on to the scarce data on the genetic basis of oscillatory neuronal dynamics. In addition to *COMT* and *BDNF* polymorphisms, many other genetic factors are likely to contribute to oscillation dynamics, as observed for the cortical structure ^96^, but this remains to be charted in future studies. Because oscillation amplitudes, LRTCs, and global synchronization dynamics are predictive of behavioral variability over both long ^36, 37^ and sub-second time scales ^4^, *COMT* Val^158^Met and *BDNF* Val^66^Met polymorphisms may underlie individual differences in trait-like behavior and cognitive performance via their effect on critical oscillatory dynamics and synchronization. Differences in these dynamics caused by genetic polymorphisms are likely to influence individuals’ cognitive and mental development and may predispose them to specific brain diseases. In line with the latter idea, alterations in the brain E/I ratio have been suggested to predispose individuals to dementia and Alzheimer’s disease ^97^. Our results thus point in new directions for the investigation of how genetic factors could translate to behavioral variability via modulations of the systems-level neuronal dynamics.

## Acknowledgments

This work was supported by the Jane and Aatos Erkko Foundation, by the Academy of Finland (SA 1266402, 1267030 to S.P., SA 1266745, 1296304 to J.M.P., and SA 1294761 to J.S.).

## Author contributions

Conceptualization, S.P. and J.M.P.; Methodology, F.S., S.P., J.M.P. and V.M.; Formal Analysis (MEG): J.S., F.S. and V.M.; Formal Analysis (Genotyping): K.K. and T.P.; Investigation: E.B.; Writing – Original Draft: J.S., S.P., F.S., J.M.P. and E.B.; Visualization: J.S., F.S. and V.M.; Funding Acquisition: E.B., J.M.P., S.P., T.P. and J.S.

## STAR Methods

### Resource availability

#### Lead contact

Further information and requests for resources should be directed to and will be fulfilled by the lead contact, Satu Palva.

#### Materials availability

Ethical restrictions apply to data and original data cannot be shared on a public server. Data underlying figures, statistics, and main conclusions will be uploaded to Dryad server upon acceptance.

#### Data and code availability

- The neuroimaging and genetic data reported in this study cannot be deposited in a public repository because ethical restrictions apply to original data. To request access, contact Satu Pal<u>va (satu.palva@hels</u>inki.fi). In addition, data underlying figures, statistics and main conclusions derived from the raw data have been deposited at Dryad server and are publicly available as of the date of publication. DOIs are listed in the key resources table.
- All original code has been deposited at https://github.com/palvalab/RS-Gen and is publicly available as of the date of publication.
- Any additional information required to reanalyze the data reported in this paper is available from the lead contact upon request.

### Experimental model and subject details

#### Human subjects

Data from 82 healthy human volunteers (age 18 to 55 years old; mean age: 29 years; 6 left- handed; 44 female) were collected for this study. This sample size corresponds to a meta-analysis of neuroimaging studies for *COMT* effects ^58^ that concluded based on pooled effect size estimates that 62 is the sample size necessary to achieve 80% power to detect association with PFC activation at an α-level of 0.05 (two-tailed). The study was performed in accordance with the Declaration of Helsinki and with permission of the Ethical Committee of the Helsinki University Central Hospital. All participants gave a written informed consent prior to the recordings.

Our sample for the *COMT* gene consisted of 18 Val/Val, 48 Val/Met, and 16 Met/Met individuals (Table 1). A one-way ANOVA confirmed that the *COMT* polymorphism groups did not differ in terms of age [*F*(2, 46) = 0.05, *p* = .984] and the duration of the resting state recording [*F*(2, 46) = 0.32, *p* = .726]. A chi-square test further confirmed that the percentages of female and male participants did not differ between the *COMT* polymorphism groups [*X^2^*(2) = 2.11, *p* = .349]. The *BDNF* sample consisted of 66 Val/Val, 14 Val/Met, and 2 Met/Met individuals (Table 2). As the Met/Met polymorphism had a low frequency, we combined Met carriers (*i.e.,* Val/Met and Met/Met polymorphisms) into one group see also ^61^. An independent samples *t*-test (2-tailed) confirmed that the *BDNF* polymorphisms did not differ in terms of age [*t*(32) = 0.20, *p* = .847] or the length of the resting state recording [*t*(32) = 0.76, *p* = .444]. A chi-square test showed no differences in the percentages of female and male participants between the *BDNF* polymorphisms [*X^2^*(1) = 0.00, *p* = .995].

#### Neuroimaging data

Eyes open resting-state brain activity was recorded from all participants with 306-channel MEG (Vectorview, Elekta-Neuromag, LtD) at a sampling rate of 600 Hz for ∼8 min (duration 7.8 ± 2.9 min, mean ± standard deviation, Tables 1 and 2). T1-weighted anatomical magnetic resonance imaging (MRI) MP-RAGE scans were obtained for each participant at a resolution of 1 × 1 × 1 mm using a 1.5 T MRI scanner (Siemens, Germany).

#### Genetic data

Blood samples for the collection of DNA were obtained from all subjects before each MEG session and DNA was extracted from blood samples according to standard procedures at the Finnish National Institute for Health and Welfare. DNA samples were genotyped at Estonian Genome Center using Infinium PsychArray-24 v1.1 (Illumina). Quality control (QC) was performed with PLINK 1.9 (www.cog-genomics.org/plink/1.9/). Markers were removed for missingness (> 5%) and Hardy-Weinberg equilibrium (*p* < 1 x 10^-6^). Individuals were checked for missing genotypes (> 5%), relatedness (identical by descent calculation, PI_HAT > 0.2) and population stratification (multidimensional scaling). The dataset collected for this study is part of a broader project “Tunteet” that includes also other datasets and paradigms.

### Method details

#### Preprocessing, filtering, and source analysis of MEG data

Signal space separation method (tSSS) ^98^ with the Maxfilter software (Elekta Neuromag, Helsinki, Finland) was used to suppress extracranial noise, interpolate bad channels and to co- localize the recordings in signal space individually for each participant. Next, independent component analysis (ICA, Matlab toolbox Fieldtrip ^99^ was used to extract and exclude signal components that were correlated with eye movement, blink and cardiac artifacts. The preprocessed MEG data were filtered into 26 logarithmically spaced frequency bands, *f_min_* = 3 Hz; *f_max_* = 60 Hz using complex Morlet Wavelets with parameter *m*=5 (making a compromise between time and frequency resolution). After filtering, the time-series data were decimated to a frequency-dependent sampling rate of between 2 and 5 times the center frequency. FreeSurfer software (http://surfer.nmr.mgh.harvard.edu/) was used for volumetric segmentation of the MRI data, surface reconstruction, flattening, cortical parcellations and neuroanatomical labeling with the Destrieux atlas. We then iteratively split the largest parcel in the atlas in two, until we arrived at a resolution of 400 parcels. MNE software (https://mne.tools/stable/index.html) ^100^ was used for creating cortically constrained source models and for the preparation of the forward and inverse operators ^101^ using noise covariance matrices (NCMs) obtained from broad-band filtered data (125–195 Hz, with 150 Hz notch). MNE inverse operators were then computed for all wavelet frequencies and used to project the sensor-space data into source-space. The source models had dipole orientations fixed to the pial surface normal, yielding 5000–8000 source vertices per hemisphere. As in previous studies ^15, 102^, source-vertex time-series were then collapsed into parcel time series with an individually source-reconstruction-accuracy- (fidelity-) optimized collapse operators.

#### Estimation of oscillation dynamics

The parcel-wise narrow-band filtered time-series were used for cortex-wide mapping of induced oscillation amplitudes (A) and LRTCs in oscillation amplitude envelopes, as well as for mapping cortex-wide phase-synchronization networks. Data were averaged for each cortical parcel and frequency separately for subjects in each *COMT* and *BDNF* polymorphisms group. Oscillation amplitudes were extracted from Morlet-wavelet filtered complex-valued time-series while LRTC exponents of the amplitude envelopes were quantified with detrended fluctuation analysis (DFA) ^70^. The power law scaling exponent β was defined as the slope of linear regression of the function *F(Δt)* plotted in log–log coordinates, estimated using a least-squares algorithm. To estimate Δ cortex-wide synchronization connectomes, phase synchronization between all 400 parcels was computed using the weighted phase-lag index (wPLI) ^15, 103^. The wPLI is insensitive to artificial connections arising as direct effects of zero-phase lagged linear signal mixing that is a major issue in connectivity analysis using MEG/EEG data ^4^. To reduce the amount of spurious interactions, we removed from the analysis parcels with *fidelity* < 0.2, which led to the exclusion of 9.27%, of possible parcel pairs ^15, 102^. The mean connection strengths were obtained by averaging the strength of all remaining edges for each subject and then over subjects in each *COMT* and *BDNF* polymorphism group. For functional subsystem analysis, all pairwise interactions were averaged across the frequencies in one of the bands defined by clustering analyses (Figure S1) and then collapsed into the 7 functional systems of the Yeo parcellation ^72^.

#### Computational modeling

We used a nested Kuramoto model ^15^ to investigate the covariance and mutual correlations of local synchronization (node/parcel amplitude), inter-areal synchronization, and local LRTCs in a heuristic model of synchronization dynamics. The model was adapted from a conventional Kuramoto model so that it comprised 100 nodes (corresponding to 100 cortical parcels) and each node was modelled by a Kuramoto model of 500 oscillators. The model was defined so that for each node *k*, the temporal evolution of the phase *θ* of each oscillator *i* was described by:

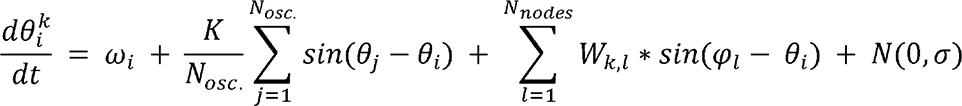

where *W_i_* is the oscillation frequency of an oscillator, K is the local coupling (control) parameter, is the average phase of the node *l*, is the connection strength between the node *k* and other nodes *j*, and *N*(0,σ), is Gaussian noise. We set all oscillators’ frequencies to a constant value (10 Hz). *W* was approximated from the log-scaled structural connectome of the Schaefer parcellation with 100 parcels ^104^ multiplied by a global coupling (control) parameter *L* at *L* = 0.5. Node time series mimicking the parcel time series in MEG data were obtained by averaging the complex time series of the oscillators in the node. Local synchronization was measured by the absolute value of the complex node time series (Kuramoto order) that is comparable to the amplitude of MEG parcel time series. We predicted the synchronization dynamics of neuronal oscillations for two hypothetical cohorts that differed in their control parameters regulating their position in the critical regime. (A, control parameter from 8.25 − 9.25; and B, control parameter from 8.75 − 9.75) operating in a subcritical regime. Because modeled data, unlike MEG data, is not influenced by volume conduction we chose to estimate pairwise synchrony here with the Phase Locking Value (PLV) which in the absence of volume conduction is roughly equivalent to wPLI, but more computationally efficient. As with MEG, we used DFA to estimate LRTC exponents. We estimated the correlation between amplitudes and DFA exponents, and between synchronization and DFA exponents, across all nodes, with Spearman’s correlation coefficient.

### Quantification and statistical analysis

Before statistical testing, time-series were collapsed into the 148 parcels of the original Destrieux atlas ^105^ to limit the number of comparisons and to improve statistical stability by averaging out anatomical variability, which is large across individuals. Group-level upper and lower confidence limits (2.5% and 97.5%) were computed for each polymorphism group with a bootstrapping approach, using *N* = 1000 resamplings with replacement of the subjects in the group.

To obtain canonical frequency bands, we first estimated spatial similarity across frequency-bands separately for oscillation amplitudes and DFA exponents using machine learning by Louvain community detection ^106^ with gamma value of 1.09. This analysis identified canonical frequency bands of theta (θ, 3–7 Hz), alpha (α, 8–14 Hz), beta (β, 14–30 Hz), and gamma (, 30–θ α β γ 60 Hz) (Figure S1). We used repeated-measures analysis of variance (ANOVA) (IBM SPSS Statistics) for testing differences in the oscillation amplitudes (A) and DFA exponents averaged across all parcels between the frequency-bands. The effects of *COMT* and *BDNF* polymorphisms on oscillation amplitudes and DFAs over the whole brain were tested separately for each Morlet, and ) using univariate γ ANOVA with the polymorphisms as between-participants factors and age as a covariate. Partial eta squared (*η* ^2^) is reported as a measure of effect size.

We then used the Mann-Whitney-U test (*p* < 0.05) to test pair-wise differences between the polymorphism groups separately for each wavelet frequency (Figure 2B, 3B) and parcel (Figure 2C, 3C). Multiple comparisons were initially accounted for with false discovery rate reduction (FDR) by discarding as many least-significant observations as were predicted to be false by the alpha-level (5% at α = 0.05) for each time-frequency bin. We further estimated a threshold (see dark grey area in Figures 2B and 3B) for the residual fraction of significants so that the probability, *p_q_*, for finding a fraction of significant value above this threshold by chance in any single frequency out of all wavelet frequencies is *p_q_* < 0.05.

For inter-areal synchronization, confidence intervals were computed with a bootstrapping approach (*N* = 1000). As the synchronization data violated the normality assumption, we tested, for each Morlet frequency, whether there was a significant difference between polymorphism groups with a nonparametric Kruskal-Wallis test (*p* < 0.05) and then used the Benjamini-Hochberg method to correct for multiple comparisons (Figure 4A). Connection density (*K*) of the mean connection strengths was used to index the proportion of significant edges from all possible interactions. Differences between the polymorphism groups in connectivity within and between subsystems ^72^ were then tested for statistical significance with the Kruskal-Wallis test (*p* < 0.05) and corrected for multiple comparisons with Benjamini-Hochberg (Figure 4B).

Finally, we estimated the correlation coefficient between amplitudes and DFA exponents and between synchronization and DFA exponents, for each parcel and wavelet frequency, across subjects within the polymorphism groups, using Spearman’s rank correlation test. Values were then averaged across all parcels and within frequency bands. Confidence limits from 2.5% to 97.5% were obtained from bootstrapping (*N* = 1000) as above. Significance testing for difference between polymorphism groups was done with permutation statistics (*N* = 1000) where the subjects were randomly assigned, without replacement, to original-sized polymorphism groups.

