## Supplemental information for "Genetic polymorphisms in *COMT* and *BDNF* influence synchronization dynamics of human neuronal oscillations"

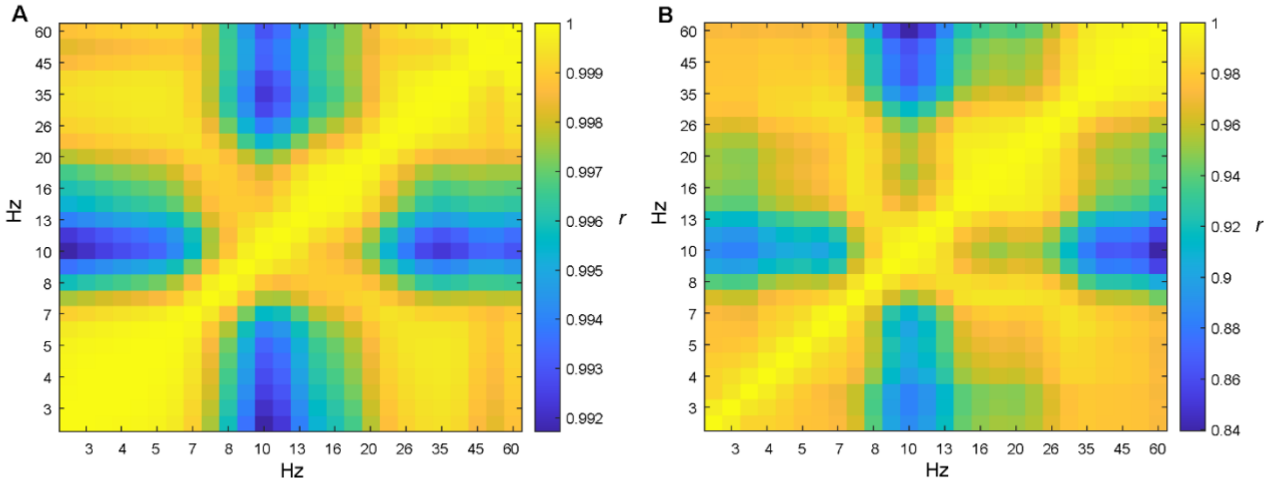


**Figure S1. Frequency-band clustering.** Spatial similarity across frequency-bands obtained by Louvain community detection [S1] for (A) oscillation amplitudes and (B) DFA exponents averaged across participants (*N* = 82). The clustering on spatial similarity yielded the frequency–frequency-band clusters of theta (θ, 3–7 Hz), alpha (α, 8–14 Hz), beta (β, 14–30 Hz), and gamma (γ, 30–60 Hz) bands. The color scales indicate the correlation coefficients.

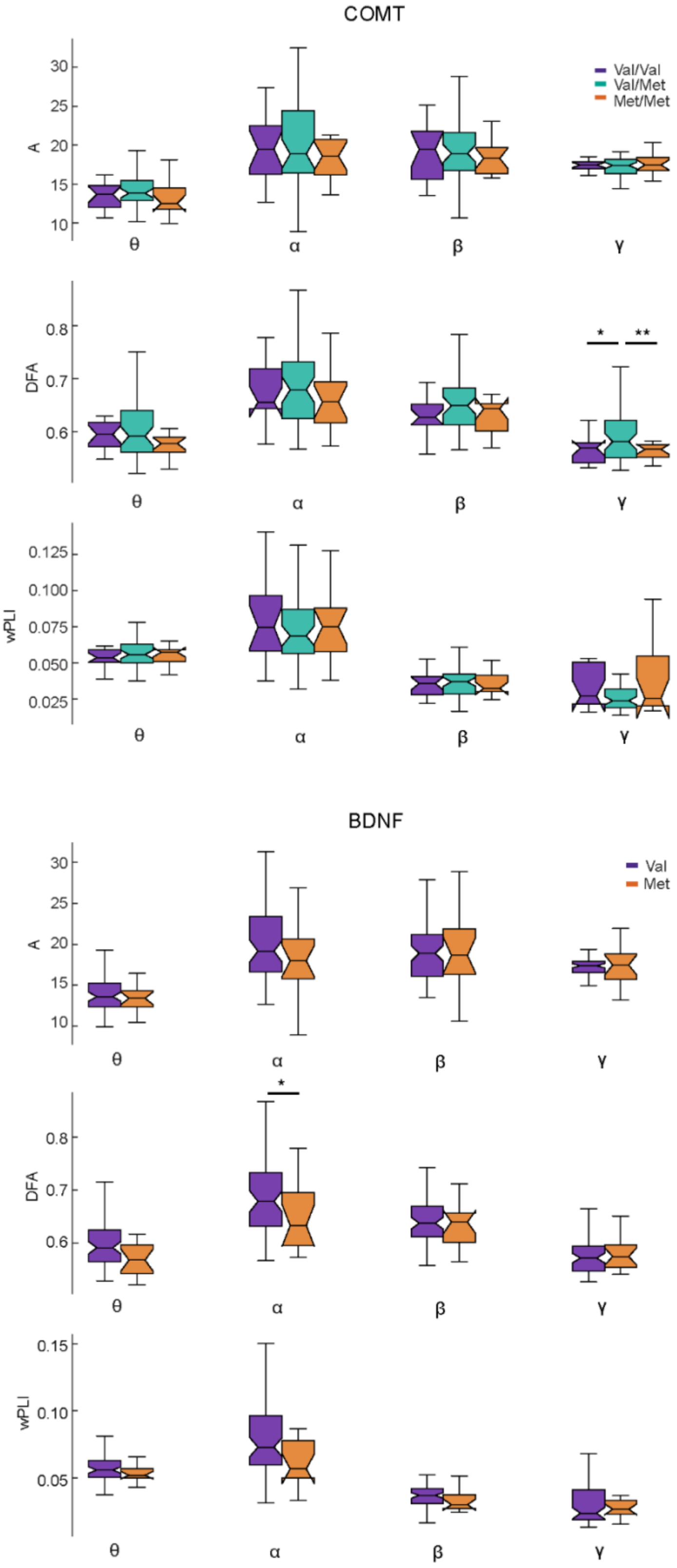
COMT
Val/Val
Val/Met
Met/Met
A
DFA
wPLI
θ
α
β
γ
*
**
BDNF
Val
Met

**Figure S2. Whole brain averaged oscillation amplitudes, DFA and phase synchronization.** Box plots of A, DFA and wPLI averaged over theta (θ, 3–7 Hz), alpha (α, 8–14 Hz), beta (β, 14–30 Hz), and gamma (γ, 30–60 Hz) frequency bands separately for each *COMT* (Val/Val in purple $n$ = 18, Val/Met in turquoise $n$ = 48, and Met/Met in orange $n$ = 16) and *BDNF* polymorphism (Val/Val in purple $n$ = 66 and combined Val/Met & Met/Met in orange $n$ = 16) groups. The box ends indicate the lower quartile (Q1, 25th percentile) and upper quartile (Q3, 75th percentile). The central notches denote the median, and the whiskers correspond to the range of Q1 – 1.5 * IQR and Q3 + 1.5 * IQR (where IQR is the inter-quartile range) and give roughly a 95% CI for comparing the medians. Asterisks denote significant group differences (*: $p < 0.05$, *: $p < 0.01$, t-test with unequal variances).

Statistical analysis using repeated measures ANOVA demonstrated a main effect of frequency band [$F$(1.86, 150.76) = 98.54, $p$ = 6.47E-27, $\eta_p^2$ = 0.549] ($N$ = 82) on oscillation amplitudes (A) at the whole-brain level. Post-hoc tests indicated that the oscillation amplitudes were stronger in the α band than in the θ ($t$(81) = 14.14, $p$ = 8.93E-23), β ($t$(81) = 2.75, $p$ =0.043), and γ ($t$(81) = 5.11, $p$ = 1.3E-5) bands (t-test, Bonferroni corrected). β-band amplitudes were also stronger than in θ ($t$(81) = 15.71, $p$ = 1.57E-25) and γ ($t$(81) = 5.49, $p$ = 3.0E-6) bands, and γ amplitudes were stronger than θ band amplitudes ($t$(81) = 12.73, $p$ = 2.75E-20).

Similar to oscillation amplitudes, repeated measures ANOVA showed a main effect of frequency for DFA exponents [$F$(2.16, 174.70) = 123.65, $p$ = 9.85E-36, $\eta_p^2$ = 0.604] ($N$ = 82). Post-hoc test indicated that DFA exponents in α band were stronger than in θ ($t$(81) = 13.00, $p$ = 1.00E-22), β ($t$(81) = 10.00, p = 1.14E-11) and γ ($t$(81) = 12.80, $p$ = 3.58E-

21) bands (t-test, Bonferroni corrected). Similarly, also β band DFA exponents were stronger than in θ ($t(81) = 10.33$, $p = 5.09E\text{-}14$) and γ ($t(81) = 10.75$, $p = 6.04E\text{-}17$) bands, and the DFA exponents in θ band were stronger than in γ band ($t(81) = 3.00$, $p = 0.008$).

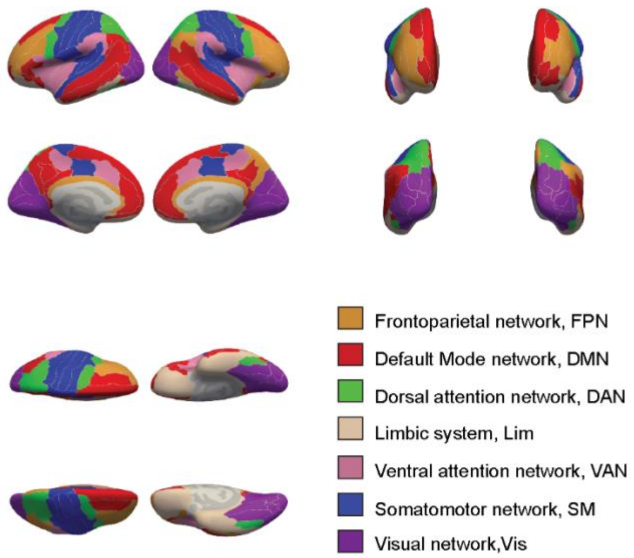


**Figure S3. The functional subsystems.** Cortical parcels of the Destrieux atlas [S2] plotted on inflated cortical surface. Color indicates the functional subsystems defined by fMRI functional connectivity [S3].

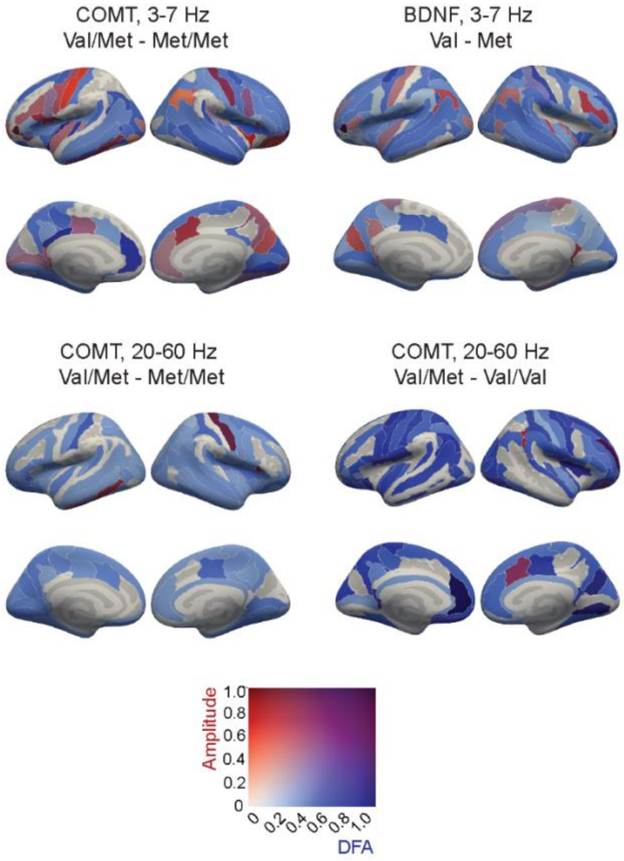


**Figure S4. Co-localization of oscillation amplitudes and LRTCs.** Cortical parcels in which amplitudes and DFA exponents were significantly larger for *COMT* Val/Met than Met/Met polymorphism groups in θ (3–7 Hz, upper left panel) and in β-γ (20–60 Hz, lower left panel) band, and for *BDNF* Val homozygotes than Met-carriers in θ (upper right panel), as well as for *COMT* Val/Met than Val/Val polymorphism groups in β-γ (lower right panel). The color indicates the difference in oscillation amplitudes (red) and DFA exponents (blue), as well as their joint effects (purple).

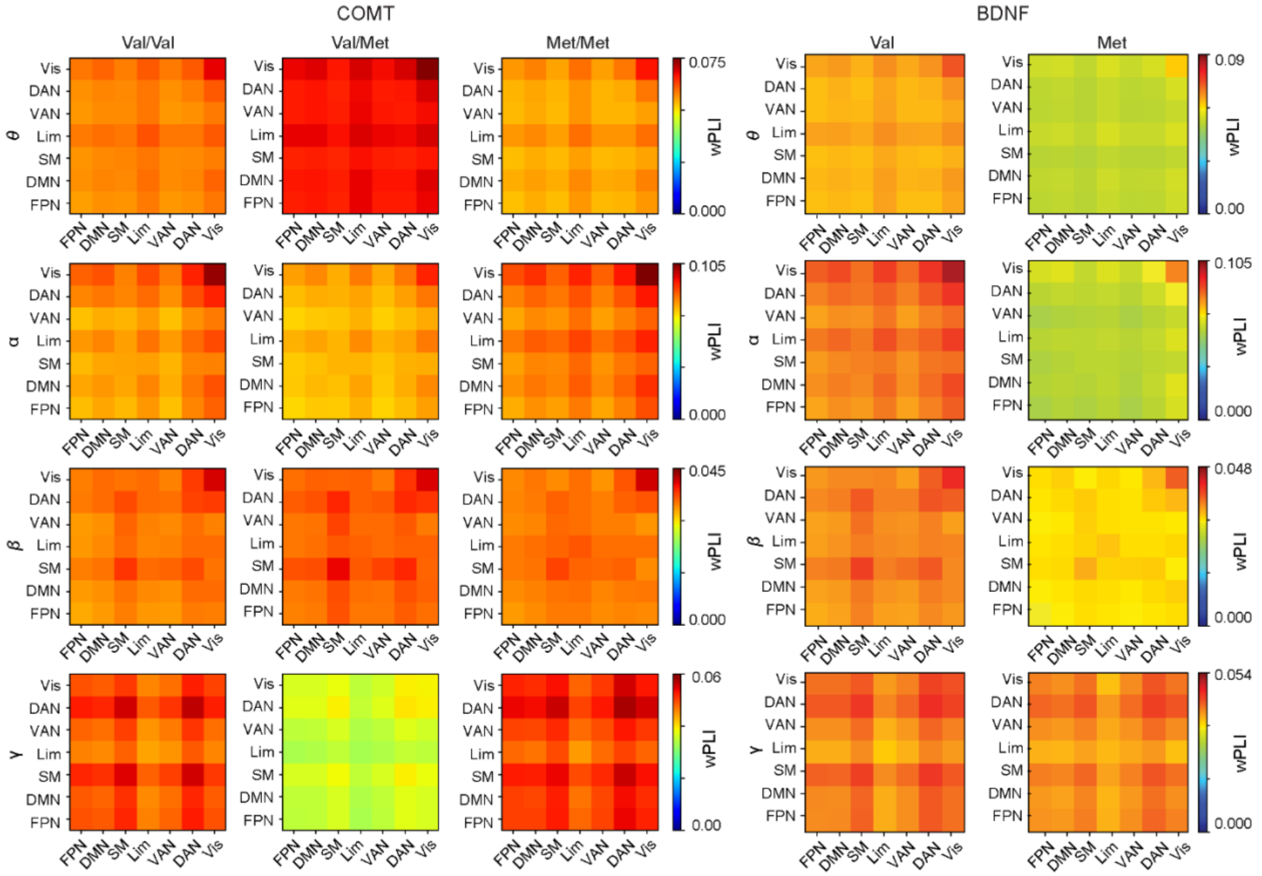


**Figure S5. Phase synchronization associated with *COMT* and *BDNF* polymorphisms.** The mean phase-synchronization within and among functional Yeo subsystems [S3] across the whole brain in the canonical frequency bands for the *COMT* Val/Val, Val/Met and Met/Met polymorphism groups and the *BDNF* Val/Val alleles and Met carriers. Inter-areal phase phase-synchrony was estimated between and within functional subsystems using the weighted phase-lag index (wPLI), of which the strength is indicated by the color scales. No differences in inter-areal phase coupling were found between *COMT* polymorphisms (left panels). *BDNF* Val/Val alleles showed stronger synchronization than Met carriers particularly in the α, β and γ bands (right panels), these differences are shown in Figure 4C. Note that the scales vary across the frequency bands. Abbreviations of the functional subsystems as in Figure S3.

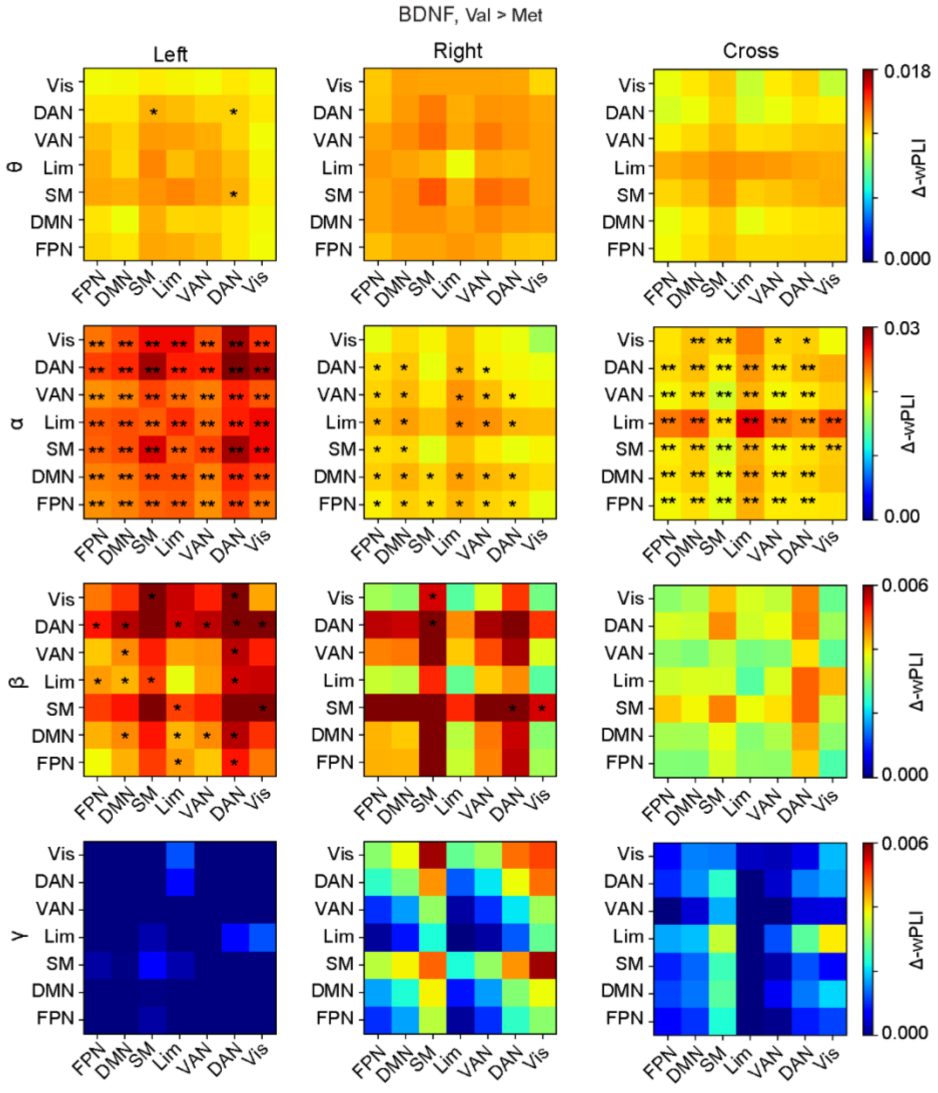


**Figure S6. Hemispheric phase synchronization for the *BDNF* polymorphism groups.** The differences in the mean phase synchronization, as estimated with wPLI, within and between functional subsystems separately for left**-**, right**-** and cross-hemispheric connections. *BDNF* Val/Val homozygotes had stronger phase synchronization compared to Met-carriers averaged over frequency bands. Color indicates the Val > Met difference in mean phase-synchronization. Stars denote network pairs where the group difference was significant (Kruskal-Wallis, *: $p < .05$**,** uncorrected,

**: $p < .01$, corrected with Benjamini-Hochberg). Abbreviations of the functional subsystems as in Figure S3.

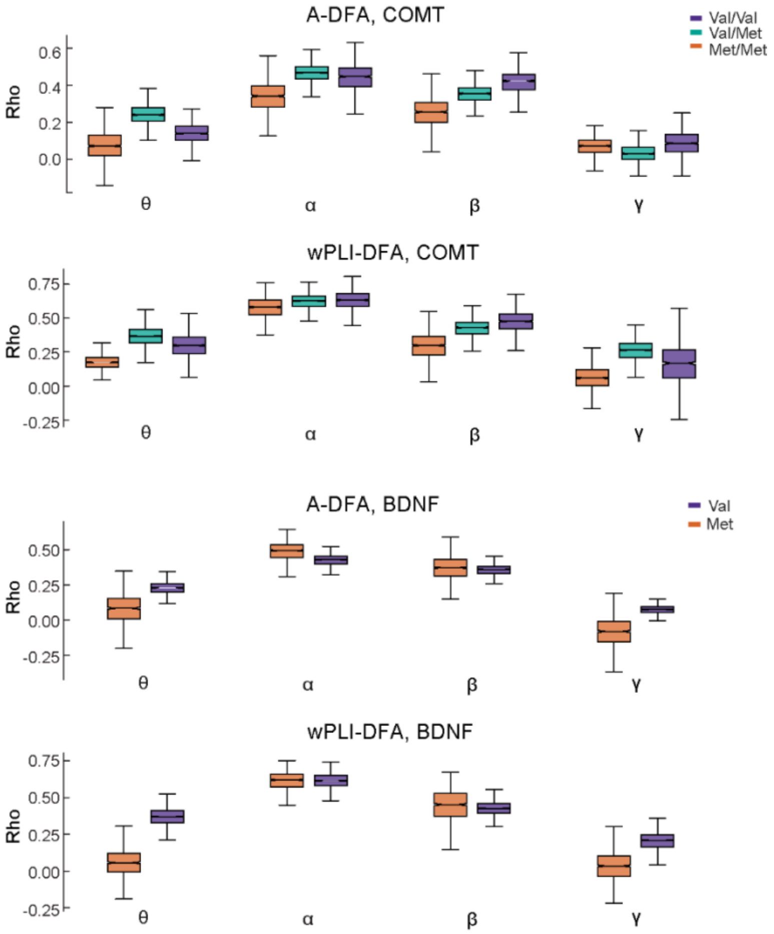


**Figure S7. Correlations of local and global synchronization with LRTCs.** Box plots of correlations of between A and wPLI with DFA exponents averaged over theta (θ, 3–7 Hz), alpha (α, 8–14 Hz), beta (β, 14–30 Hz), and gamma (*γ,* 30–60 Hz) frequency bands. Correlations are shown separately for each *COMT* (Val/Val, Val/Met, and Met/Met, upper panels) and each *BDNF* (Val/Val and Met-carriers, lower panels) polymorphism group. Box plot details as in Figure S2.
